## Supplemental File 1 for "Living apart together: modelling the consequences of the dikaryotic life cycle of mushroom-forming fungi for genomic conflict"

read input parameters from Temporary_Input.dat in same folder

##!Initial State##
for each i in initial_spores:
   choose random x and y
   if empty (i.e. mating type == 0)
   place spore of random mating type
   gaussian random assign fitnesses
  

##!Site Update##
do update for (grid_size * grid_size) times
   pick random x and y
   if empty (i.e. mating type == 0) 
      skip death routine
   else
      pick random number
      if less than death rate
         kill cell, remove mating type and fitnesses
         end update
   ##attempt sex in female role##
   if site is monokaryotic, or if dikaryotic and is open dikaryon
      initiate mating
      if no mates found
         end update
      spread nuclei of mating throughout individual of focal female
      if the mating partner was a dikaryon and open dikaryon
         spread nuclei of mating throughout individial of mating male as well
      end update
   #vegetative growth
   get random number
   if lower than basal growth rate
      get fitnesses of 4 neighbours
      the best neighbour grows in, ties are broken by random
      end update
   #spore disperal
   for each site in grid
      iterate for number of spores in basal spore production
         produce spores if fitness is higher than random number
         place spore randomly based on dispersal
   when finished, spores land if site is empty, if multiple spores the winner is random
iteration is finished, calculate statistics and perform next iteration
      
