## Supplementary figures and images for "Living apart together: modelling the consequences of the dikaryotic life cycle of mushroom-forming fungi for genomic conflict"

### Supplemental File 2: Diploid animation

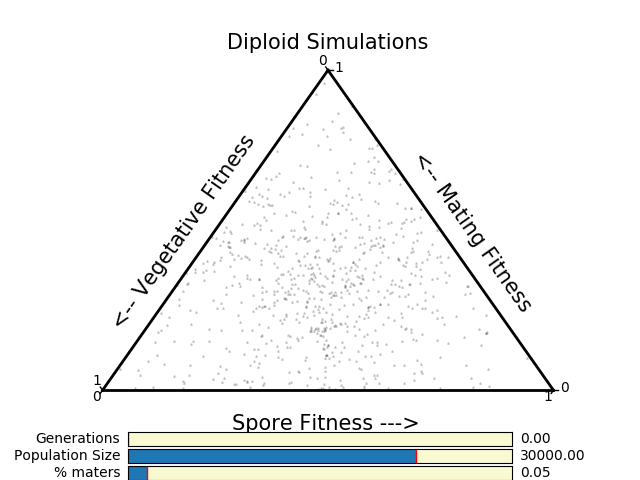

### Supplemental File 3: Standard animation

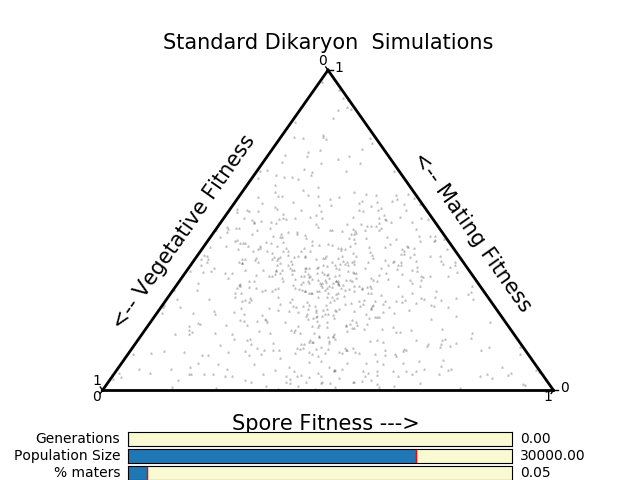

### Supplemental File 4: Open dikaryon animation

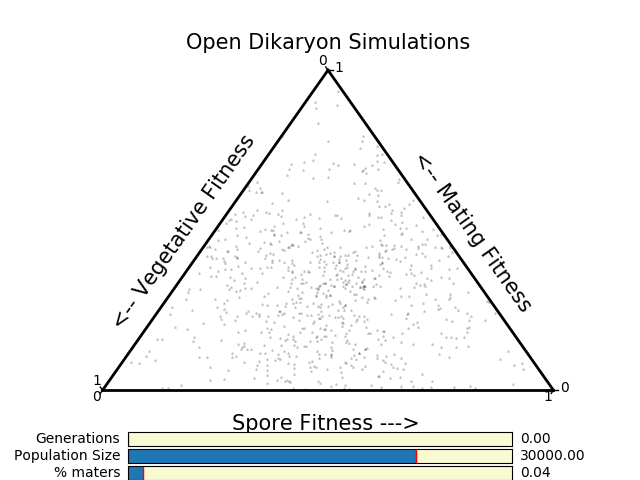
